# Proteome-wide discovery of novel diagnostic antigens for coccidioidomycosis

**DOI:** 10.64898/2026.11.05.764335

**Authors:** Evan A. Elko, Luke Ryle, Georgia A. Nelson, Nathan G. Sarbo, Caroline E. Harms, Erin Kelley, Megan N. Fredericks, Paul Keim, Kenneth Knox, Mrinalini Kala, Deborah H. Fuller, Erik W. Settles, Bridget M. Barker, D. Mitchell Magee, Douglas F. Lake, Thomas E. Grys, George R. Thompson, John A. Altin, Jason T Ladner, Heather L. Mead

## Abstract

Coccidioidomycosis, commonly called Valley Fever (VF), caused by *Coccidioides* spp., is an understudied fungal disease that accounts for a substantial fraction of pneumonias in endemic areas. Diagnosis remains challenging because the mainstay of diagnosis is serology, and current assays have modest sensitivity in early infection. Discovery of new antigens could improve serologic detection early in disease. Here, we use PepSeq, a DNA-programmable serology platform, to represent the complete proteome of *Coccidioides* within a library of 244,000 DNA-barcoded peptides. By profiling antibody reactivity at the protein and epitope levels in VF patients, naturally infected pigtail macaques, and healthy controls, we identify numerous previously-unrecognized antigens and epitopes, including some shared between humans and macaque responses. When translated into a pooled peptide ELISA, we show that a subset of identified antigens can discriminate VF cases from controls in an independent cohort with high accuracy (AUC = 0.91). These findings establish high-dimensional peptide serology as a scalable approach for fungal antigen discovery and identify new targets for improved VF diagnosis.

## Introduction

Coccidioidomycosis, commonly called Valley Fever, (VF) is an environmentally acquired fungal disease endemic in the southwestern United States, where its incidence has been increasing in recent years^1^. The disease has a broad range of clinical manifestations, ranging from mild and self-limiting infections to chronic pneumonia^1^. In rare cases, the infection can become extrapulmonary and spread to other organs^2,3^. Due to the presentation of symptoms similar to those of many bacterial infections, delays in diagnosis are common, often resulting in inappropriate use of antibiotics and increased risk of developing severe disease caused by the delay in correct evaluation and treatment^3,4^. Even in endemic areas where awareness is high, a correct medical diagnosis typically requires multiple clinical visits and takes a median of 23 days^4^.

Clinicians frequently rely on serology-based assays to aid in diagnosis of VF, as they are rapid, more automatable, and are technically less complex than culture or microscopy based methods^2,5^. All available methods have important limitations in terms of turnaround time, clinical performance, and/or technical complexity and thus there is not a standard method rather clinicians rely on a combination of serology tests^2,6^. Current serological diagnostics typically use crude fungal lysate to detect responses against one of three known diagnostic antigens: Beta-glucosidase 2 (formerly referred to as tube precipitin, TP)^7,8^, complement fixation chitinase (CTS1, CF)^9^ and galactomannan^10^. These targets have been used in several different formats, including enzyme immunoassay (EIA)^11,12^, lateral flow^13^, immunodiffusion, and complement fixation^14^, where they primarily detect either IgM or IgG reactivity^6,11^. The sensitivity of these assays range from ~30-87%, with specificity ranging from ~68-99%.^5,15–18^. Considerable effort has been put into increasing the performance of these assays and identifying new antigenic targets, however, identifying and testing potential antigens has been challenging due to the large number of potential immunogenic proteins.

*Coccidioides* spp. are genetically and morphologically complex, with ~8,000 protein coding genes per strain, 50% of which are unannotated^19–21^. Identification and production of unannotated proteins with immunogenic potential is labor intensive (crude lysate, SDS page, sequencing) and cost prohibitive (full length protein expression and purification at scale). Additionally, the number of antigens (once identified and purified) that could be tested for antibody reactivity has been limited by the cost and sample volume requirements associated with establishing and running single-plex serology assays, such as enzyme-linked immunosorbent assays (ELISAs)^22^. However, recent advances in highly-multiplexed serology now allow for bulk antigen synthesis and simultaneous assessment of antibody binding for hundreds of thousands of antigens with minimal upfront investment and sample consumption^23,24^. Which allows for full proteome interrogations of complex pathogens, like *Coccidioides*.

PepSeq is a fully *in vitro* and highly-multiplexed serology platform that generates and uses custom libraries of DNA barcoded peptides to quantify antibody responses with epitope-level resolution^23^. We have previously used this technology to characterize antibody responses to large numbers of viruses and bacteria at proteome-wide scales and through the lenses of natural infection and vaccination^25–29^. Here, we created a novel 244,000-member peptide library, *Coccidioides* whole proteome (CWP), which was used to assess reactivity profiles for IgG, IgA and IgM antibodies in serum samples from human subjects with confirmed VF or blood bank donor controls, as well as IgG profiles for plasma samples from naturally infected pigtail macaques (PTM). We next created a novel multi-peptide ELISA based on the significantly reactive peptides identified in our IgG screen and were able to discriminate between human VF and control patient sera in a validation cohort. Thus, demonstrating that the epitopes discovered using PepSeq may be useful for the improvement of VF diagnostic approaches.

## Methods

### Serum/Plasma samples

The discovery VF cohort (n=70) came from the University of Arizona biorepository. The discovery controls ( n=60) were treated as presumed negative controls as they were collected from blood bank donors outside the endemic range for VF (Louisiana, USA). The Discovery cohort were also assayed on the ELISA but as a distinct group. The ELISA validation VF cohort was obtained from multiple sources: the UC Davis Coccidioidomycosis Serology Laboratory (n= 16), the University of Arizona biobank (n=2) and the Mayo Clinic (n=60). The ELISA validation negatives were divided into two groups 1) additional presumed negative controls (n=30) collected from blood bank donors outside the endemic range for VF (Louisiana, USA) and 2) endemic negative controls (n=32) was provided by the Mayo Clinic.

Plasma samples were obtained from naturally infected pigtail macaques (n=33) at the Washington National Primate Laboratory (WaNPRC). Animals at the WaNPRC Arizona Breeding Colony are housed in Mesa, AZ. As a part of routine animal care, animals undergo semi-annual exams, which include the collection of sera to test for *Coccidioides* spp. seropositivity. Excess sera from these exams are stored in a repository at WaNPRC and were shared for these experiments. Plasma samples from WaNPRC PTM animals born and raised in Seattle, WA (n=17) were used as the negative controls.

University of Arizona biorepository collected samples under IRB number, 1712117680A003, and the use of these samples for this study was approved under IRB number STUDY00002075. UC Davis Coccidioidomycosis Serology Laboratory collected samples under IRB number 1661992. The Mayo Clinic collected samples under IRB 12-000965. The Washington National Primate Laboratory collected samples under IACUC 4551-05 and IACUC 4551-04.

### Design of Coccidioides whole proteome peptide library

For the CWP PepSeq library, we targeted all predicted protein sequences for each *Coccidioides* species: *C. immitis* and *C. posadasii*. Specifically, we included all the proteins in the two publicly available genomes with the highest sequencing depth for each *Coccidioides* species, as well as a few additional sequences for known *Coccidioides* spp. antigens. For *C. posadasii*, we included Silveira 2022 (GCA_018416015.2) and RMSCC 3488 (GCA_000150055.1); and for *C. immitis*, we included WA_211 (GCA_004115165.2) and RMSCC 2394 (GCA_000149895.1). Predicted protein sequences from these four *Coccidioides* spp. genomes were downloaded from GenBank on July 20, 2023 (Supplemental Table 1). We confirmed the inclusion of all known *Coccidioides* spp. antigen protein sequences from previously reported studies^21^, Immune Epitope Database (IEDB)^30^ and the currently used diagnostic proteins.

**Table 1.** Protein characteristics for the six significantly elevated proteins in VF patient samples.

| Protein Name | Protein Accession | Localizations | Signals | Membrane types | Transcriptomic Data | Potential annotation |
|---|---|---|---|---|---|---|
| SOWgp | TPX21266.1, QVM09276.1 | Extracellular |  | Lipid anchor\|Soluble | upregulated in spherules | spherule outer wall glycoprotein |
| IgG antigen 1 | KMP05713.1, KMM71006.1, TPX21976.1 | Extracellular | Signal peptide | Soluble | upregulated in spherules | filamentous hemagglutinin |
| IgG antigen 2 | KMP00029.1, TPX26831.1 | Extracellular,Extracellular | Signal peptide, None | Soluble, Lipid anchor | upregulated in spherules | hypothetical protein |
| IgG antigen 3 | CPSG_07735 | Extracellular |  | Lipid anchor\|Soluble | upregulated in spherules | extracellular membrane protein CFEM domain-containing protein |
| IgG antigen 4 | QVM12318.1 | Cytoplasm |  | Soluble | stable expersion | geranylgeranyltransferase |
| IgG antigen 5 | TPX22474.1 | Nucleus | Nuclear localization signal | Soluble | downregulated in spherules | Zn(2)-C6 fungal-type domain |

Peptides were designed from these target proteins using a combined set cover and sliding window approach^23^. First, we generated clusters of similar protein sequences from the four focal genomes using USEARCH (70% identity threshold)^31^. We then selected representative peptides for each cluster, each of which was 30 amino acids (AA) long (https://github.com/LadnerLab/Library-Design/tree/master/SW_SC). Collectively, these peptides covered ≥95% of the unique 9mers (proxy for linear epitopes) contained in the proteomes of these four genomes and manually curated known antigens described above. Next, we removed redundant peptides using a custom script (https://github.com/LadnerLab/Library-Design/blob/master/extensions/kmerRedundancy.py) that steps through kmer sizes from 30 AA to 8 AA and removes any peptides that do not contain any unique kmers. This resulted in a peptide library containing 243,135 peptides designed from *C. immitis* and *C. posadasii* proteins. Our final design covered 96% of the unique AA 9mers in all 11 of the publicly available *Coccidioides* spp. genomes. Finally, we added 865 positive control viral peptides (designed from 40 different virus species), which previous studies have shown to be commonly recognized by IgG antibodies in human samples (Supplemental Table 2)^24,27^.

### PepSeq library synthesis and antibody binding assays

The CWP library was encoded as 244,000 128 base pair DNA oligonucleotides using custom scripts^23^, and these oligonucleotides were sourced from Agilent Technologies (SurePrint oligo pool; Santa Clara, CA) (Supplemental Table 2). These oligonucleotides were used as the starting material to synthesize the CWP PepSeq library of DNA-barcoded peptides following methods previously described^23^. Briefly, the oligonucleotide library was PCR-amplified and then used to generate mRNA in an *in vitro* transcription reaction. The mRNA products were ligated to hairpin oligonucleotide adapters bearing a puromycin molecule tethered by a PEG spacer and used as templates in an *in vitro* translation reaction, resulting in peptides covalently bound to their associated mRNA. Finally, a reverse transcription reaction, primed by the adaptor hairpin, was used to generate a cDNA tag, and the original mRNA was removed using RNAse. To perform serological assays, the resulting CWP DNA-barcoded peptide library was added to diluted plasma or serum and incubated overnight. The binding reaction was then applied to pre-washed isotype specific (IgG, IgM or IgA) capture beads. IgG was captured with streptococcal protein G-bearing beads (ThermoFisher Cat# 10004D). IgA and IgM were captured with polyclonal antibodies specific for these human isotypes. IgA and IgM specific beads were generated by incubating biotinylated anti-IgM (Biolegend #314504) or anti-IgA (Invitrogen,#MA5-27919) antibodies in excess with streptavidin beads (Thermo Fisher, #6502). Following capture, bead bound antibodies were repeatedly washed, bound PepSeq probes were eluted, and the associated DNA barcodes were amplified and indexed for sequencing. Following PCR cleanup, products were pooled, quantified, and sequenced using an Illumina NextSeq or NovaSeq instrument. All samples were assayed in duplicate.

### PepSeq Data analysis

To identify peptides with diagnostic potential for VF, we compared antibody reactivity profiles for 70 confirmed VF patients and 60 presumed negative blood bank controls (discovery cohort). To avoid bias, we focused on the subset of peptides (n=884) that showed reactivity in at least five samples, regardless of whether they were VF positive or presumed negative (Z score ≥10 in ≥5 samples). For each, a peptide-specific Z score threshold was calculated as three standard deviations above the mean Z score of the discovery non-endemic negative control samples. These peptide-specific thresholds were used to count the number of positive and negative samples in each cohort. To identify peptides with significantly more positives among our VF cohort, we used two-sided Fisher’s Exact tests with a Benjamini-Hochberg multiple test correction. Peptides of interest were identified as those with a corrected p-value ≤ 0.05. To assess our false discovery rate, we followed an identical procedure to also identify peptides with higher reactivity in 1) our control cohort compared to our VF cohort and 2) datasets with randomly permuted cohort assignments (n=1000). For each protein that contained at least one significant peptide, we predicted the protein’s cellular localization and the potential presence of a secretion signal peptide using DeepLoc2.1^32^ and SignalP-6.0^33^ with default parameters. For each protein, we input the sequence accession identifier that was used during the design process. The FungiDB database was used to search for gene/protein names, predicted function and gene expression data^34^. Sequence accession numbers were also input to NCBI BLASTP with clusterNR as the database to look for regions of similarity^35^.

To assess the diagnostic performance, within our discovery cohort, of the peptides of interest detected in our IgG screen, we generated receiver operating characteristic (ROC) curves, both for individual peptides and the summed Z scores across multiple peptides, with Z score thresholds ranging from −3.1 to 5,181.

To identify significant differences in the summed Z scores for our 31 IgG peptides of interest and CF titer, while controlling for gender, diagnosis, sample location, year of diagnosis and viral peptide counts, we utilized a generalized linear model (GLM) implemented in Python using statsmodels v0.14.1^36^. Specifically, we fit a Gaussian GLM with a single dependent variable (summed Z score) and six independent variables (CF titer [continuous], sex [categorical], diagnosis [categorical], year of diagnosis [continuous], shipment [categorical], and enriched viral peptide counts [continuous]).

### Multi-peptide ELISA’s

From an initial PepSeq screen of n=101 human samples, a set of 30 peptides of interest were identified and clustered into 11 groups based on sequence homology using Clustal Omega^37^. For clusters containing more than one sequence, the sequence with the highest Z score across all VF samples in each cluster was selected, if the cluster contained multiple potential epitopes, the peptide with the highest Z score in VF samples for each epitope was selected, resulting in 16 peptides being selected for inclusion in a novel, multi-epitope ELISA. C-terminus biotinylated, 30 amino acid peptides were obtained from Thermo Fisher (see Supplemental Table 3), and these were used to create a peptide pool at equimolar concentrations.

With this peptide pool as the antigen, a direct ELISA protocol was performed using the following materials and steps. NeutrAvidin coated plates (Thermo) were incubated at 4°C overnight with 1µg/mL of the peptide pool. Wells were blocked using a 1% Blocker BSA solution (Thermo) prior to sample introduction. Diluted (1:500) patient sample was added to each peptide containing well (Sample OD). Additionally, for each donor, diluted (1:500) patient sample was added to control wells without peptide (Serum OD). All samples and controls were run in duplicate and, for each plate, two wells with 1µg/mL of the peptide pool but without serum were included (Peptide OD). Goat anti-Human IgG-HRP conjugated antibody (Abcam) was used as the detection antibody, followed by 50µL TMB substrate solution (Thermo) for detection. Optical density was measured at 450 nm on a BioRad xMark spectrophotometer. Washes were performed between each step using a BioRad ImmunoWash 1575. The normalized OD was calculated with the following equation: Mean Sample OD - (Mean Peptide OD + Mean Serum OD). To assess the performance of the peptide ELISA, ROC curves were generated with normalized OD thresholds ranging from −0.273 to 2.527. ELISA assay was performed on a subset of the discovery cohort (51 confirmed VF patients and 55 blood bank controls) and the validation cohort, which included an independent VF sample set (n=78), additional presumed negative, blood bank controls (n=30) and endemic VF negative controls (n=32) that had not been assayed with the CWP PepSeq library. VF samples and non-endemic control samples were run side by side on the same 96-well plate to avoid any batch effects. Each 96-well plate also included a positive control and two negative control samples to help identify any plate to plate variations.

### Immunodiffusion

Three blood bank samples demonstrated reactivity to >2 of the 31 peptides of interest identified in this study. To explore whether this reactivity could be explained by past fungal infections, these serum samples were independently tested for reactivity to *Coccidioides* spp., *Histoplasma, Aspergillus* and *Blastomyces* antigens using the IMMY ID Fungal Antibody System (ID1001) following the manufacturer’s protocol. As internal controls for the assay, three VF patients from the validation cohort were also tested using the same approach.

## Results

### Identification of antibody epitopes commonly targeted in VF patients

We used the CWP PepSeq library to measure the anti-*Coccidioides* spp. IgG antibody reactivity profiles in our discovery cohort of 70 VF patients and 60 non endemic control samples (blood donors collected in Louisiana, USA - outside of the endemic range). Consistent with our expectations, both cohorts showed strong IgG antibody reactivity against a panel of 865 virus derived control peptides (93-628 peptides with Z ≥ 10 in both replicates). However, on average, we observed lower numbers of enriched viral peptides in our VF samples compared to the non-endemic control samples (median=355.5 and 431.5, respectively; p=0.001, t-test) (Supplemental Figure 1A). In contrast, we observed similar numbers of *Coccidioides* spp. derived IgG enriched peptides in VF patients and non-endemic controls (median=54 and 49, respectively; pval=0.93, t-test) (Supplemental Figure 1A).

**Figure 1.**
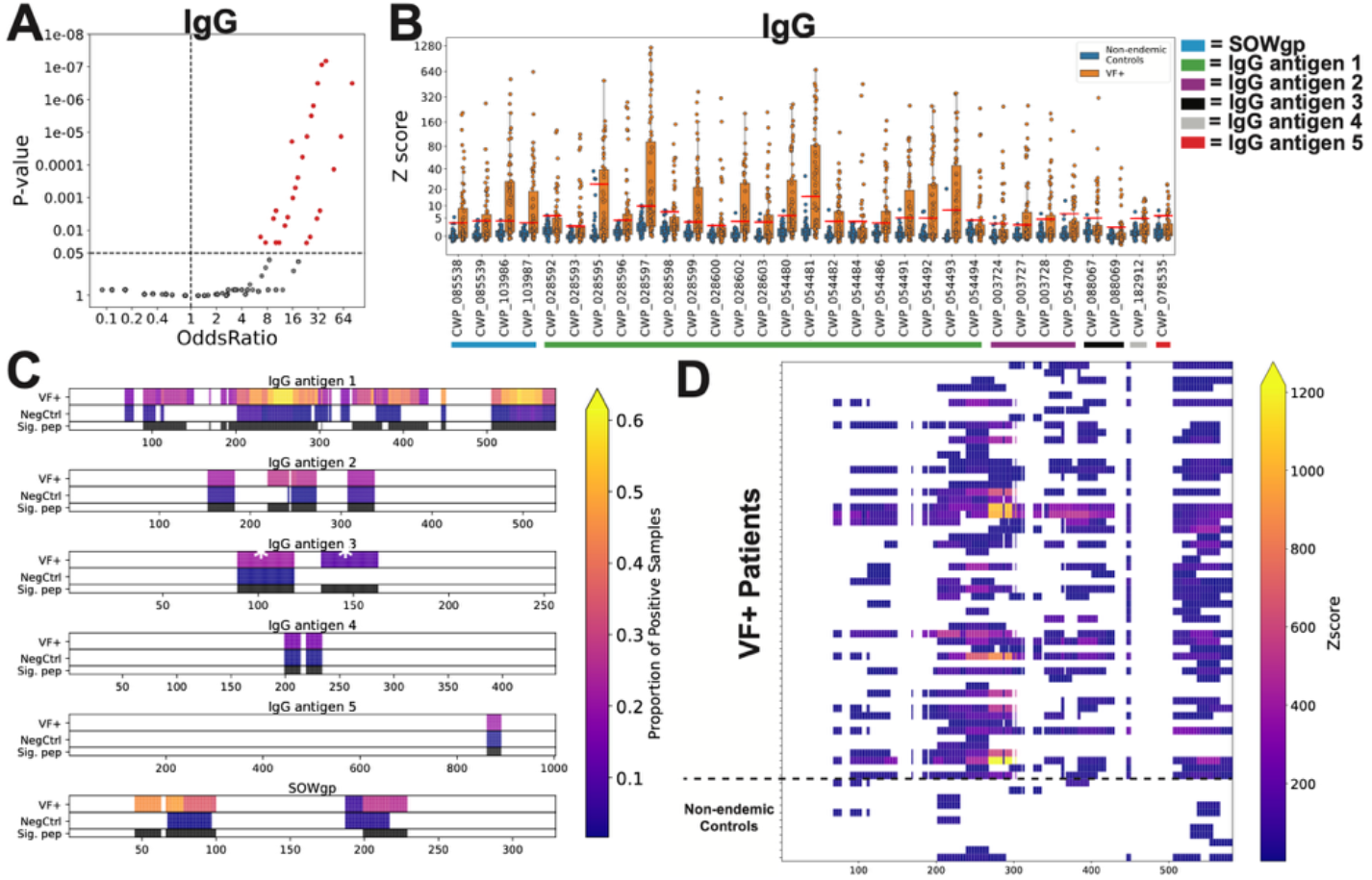
PepSeq identifies *Coccidioides* spp. peptides and proteins commonly recognized by antibodies in infected individuals. A PepSeq library (CWP) covering the full proteome of *Coccidioides* spp. was used to broadly assess anti-*Coccidioides* spp. antibody reactivity in the discovery cohort of 70 VF (Valley Fever) patient sera and 60 control sera from a non-endemic region in the US. **A)** A volcano plot showing the Benjamini-Hochberg corrected p-values from a Fisher’s exact test comparing the number of positive and negative VF and non-endemic control samples (y-axis) versus Odds Ratio (x-axis) for a subset of 884 peptides recognized by serum IgG antibodies in ≥5 serum samples. Significant peptides after multiple test corrections are highlighted in red, non-significant peptides are shown in grey. Horizontal dashed line indicates a corrected p-value of 0.05. Vertical dashed line indicates an odds ratio of 1. **B)** Boxplots showing the Z score distributions for the 31 significantly enriched IgG peptides in the VF (orange) and non-endemic control (blue). Red lines indicate the threshold set for each peptide (three standard deviations above the mean of the non-endemic control cohort). Colored bars at bottom indicate the ID of the protein from which each peptide was designed. Peptide sequences can be found in Supplemental Table 2. **C)** Heatmap showing the proportion of positive samples in both VF and non-endemic control samples across the full protein sequence for the six proteins that contain significantly enriched peptides. Black bars in “Sig. pep” row indicates the location of the 31 significantly enriched peptides identified in **A**. Asterisks in IgG antigen 3 indicate the two peptides of interest also detected in the PTM IgG screen. **D)** Heatmap showing the Z scores for reactive peptides across the IgG antigen 1 protein sequence for all samples that contained at least one reactive peptide.

To exclude peptides with little to no evidence for antibody binding, we restricted our downstream analyses to peptides that had a Z score ≥ 10 in at least 5 of the 130 total discovery samples. For the IgG assay, 884 peptides met this threshold. From our human IgG reactivity profiles, we identified 31 peptides that were significantly enriched in the VF cohort compared to our controls, after multiple test correction (Figure 1A,B), with corrected p-values as low as 6.54×10^−8^. To assess the likelihood of false positives with our analysis approach, we randomly permuted the sample names, which resulted in an average of 0.042 significantly enriched peptides per permutation, with 99.6% of permutations having no significantly elevated peptides. We also ran a flipped analysis to determine whether any of the *Coccidioides* spp. derived peptides were significantly enriched in the control cohort and found zero elevated peptides. Z score analysis showed that 87% of VF samples were reactive to at least one of the 31 identified peptides, and each VF sample, on average, was reactive to 11/31 peptides (35%) (Figure 1B). These peptides were designed from six different proteins (Table 1), with four of the six proteins having multiple significantly elevated peptides. Four of the identified peptides were designed from the known VF antigen, SOWgp (CWP_085538, CWP_085539, CWP_103986, CWP_103987) (Figure 1B). However, the remaining five proteins have not previously been described as immunoreactive. Strikingly, one of these five, which we have named IgG antigen 1, contained 19 significantly enriched peptides. Next, we considered the location of reactivity across the protein sequences. The proportion of VF patients which recognized a given epitope ranged from 0.10 to 0.63 across the six proteins (Figure 1C). For some proteins there are a small proportion (0.02-0.10) of non-endemic control samples that also showed reactivity, potentially indicating cross reactivity with other pathogens. The protein with the largest number of enriched peptides (IgG antigen 1) showed at least four epitopes that were broadly reactive across >35% of reactive VF samples, with the region from amino acid residues 200-300 showing the highest Z scores (Figure 1D).

Next, we used analogous experimental and analytical methods (only substituting a different isotype capture) to assess the IgM reactivity profiles on a subset of our discovery cohort VF patients (n=46) and non-endemic controls (n=39). There were few enriched viral peptides detected with the IgM screen in both the VF and control cohorts (median=1 and 2, respectively), and there was no significant difference between the number of *Coccidioides* spp. derived enriched peptides between the cohorts (pval=0.54, t-test) (Supplemental Figure 1B). We observed a total of 273 peptides that passed our reactivity threshold (Z score ≥ 10 in ≥ 5 samples) but found none of these were significantly enriched in the VF cohort after multiple test correction. (Supplemental Figure 2A).

**Figure 2.**
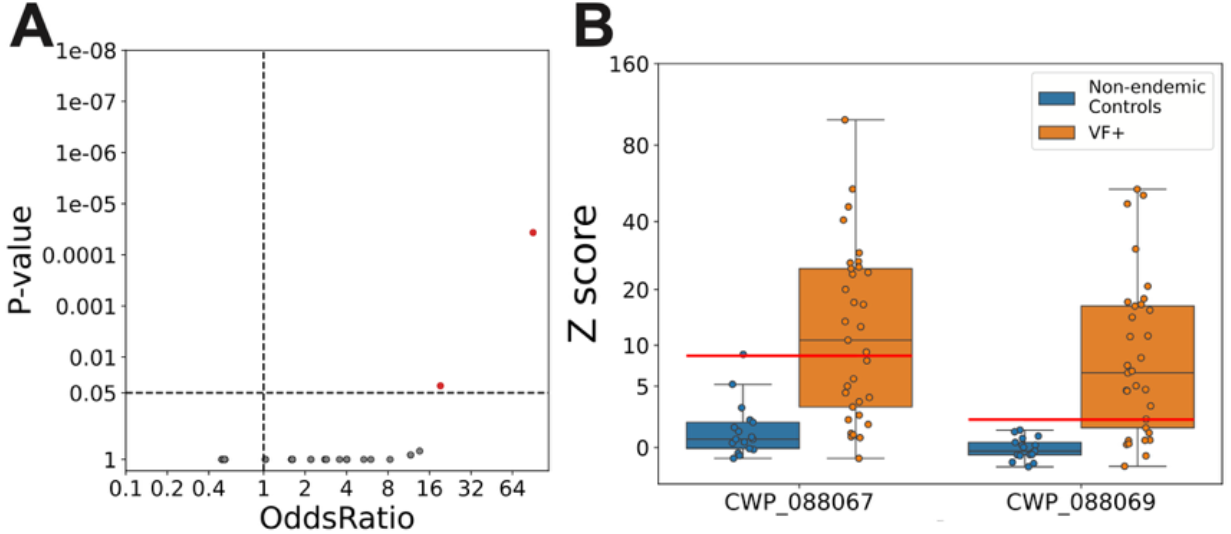
PepSeq identifies candidate diagnostic peptides with immunoreactivity in pigtail macaque serum. A PepSeq library (CWP) covering the full proteome of *Coccidioides* spp. was used to broadly assess anti-*Coccidioides* spp. antibody reactivity in the serum from 33 PTMs with confirmed natural *Coccidioides* spp. infections and 17 control sera from PTMs raised outside of the endemic region of the US. **A)** A volcano plot showing the Benjamini-Hochberg corrected p-values from a Fisher’s exact test comparing the number of positive and negative VF and non-endemic control samples (y-axis) versus Odds Ratio (x-axis) for a subset of 104 peptides recognized by PTM serum IgG antibodies in ≥5 serum samples. Significant peptides after multiple test corrections are shown in red, non-significant peptides are shown in grey. Horizontal dashed line indicates a corrected p-value of 0.05. Vertical dashed line indicates an odds ratio of 1. **B)** Boxplots showing the Z score distribution of the two significantly enriched IgG peptides in naturally infected VF PTM (orange) and the PTM raised outside of the endemic region (blue) cohorts.

Lastly, we analyzed the IgA reactivity profiles of a subset of our discovery cohort VF patients (n=50) and non-endemic controls (n=53). On average we observed a lower number of total enriched peptides (Z ≥ 10 in both replicates) in our IgA screen compared to IgG (IgA median=42, IgG median=475.5) (Supplemental Figure 1C). The number of IgA enriched viral control peptides was similar between VF and control samples (medians=24.5 and 25, respectively), however we did see a trend towards an increased number of enriched *Coccidioides* spp. derived peptides in the VF samples compared to controls (medians=8 and 5, respectively, p=0.11, t-test) (Supplemental Figure 1C). There were 32 peptides that showed reactivity in at least five samples, and two peptides that were significantly enriched in our VF cohort after multiple test correction. Both of these peptides were also detected as significantly enriched in the IgG screen and both were designed from the same protein, IgG antigen 1, the protein with the highest number of peptides with diagnostic potential in our IgG analysis (Supplemental Figure 2B,C).

### Identification of antibody epitopes commonly targeted in naturally infected pigtail macaques

Pigtail macaques (PTM) living in a captive research colony located in the endemic region (Mesa, AZ) can be naturally infected with *Coccidioides* spp^38^. Previous characterization shows that these PTMs have similar disease manifestation as seen in humans^39^. We measured the IgG reactivity profiles from PTMs with confirmed *Coccidioides* spp. infections (n=33) and control samples from PTMs housed in a separate colony outside of the endemic region (Seattle, WA) (n=17). We observed 104 peptides whose enrichment by IgG exceeded our threshold (Z score ≥ 10) in at least 5 samples regardless of disease state. We detected two significantly elevated peptides in the *Coccidioides* spp. infected PTM group, which were both designed from a single protein (IgG antigen 3) (Figure 2A,B). Both peptides were also detected as significantly elevated in the human IgG analysis (Figure 2B, Supplemental Figure 3).

**Figure 3.**
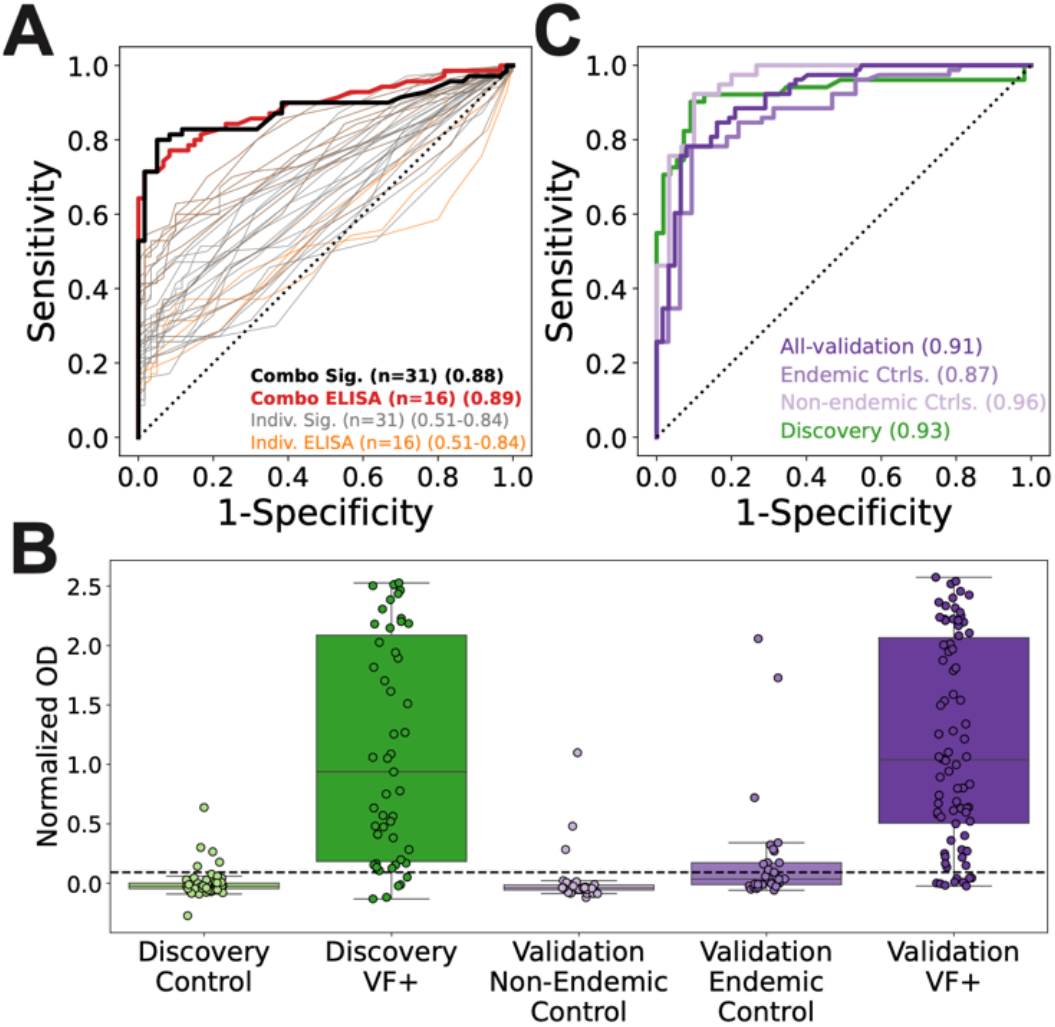
Peptides identified by PepSeq effectively discriminate between non-endemic controls and VF patients in the discovery cohort, as well as an independent VF cohort. **A)** ROC curves for discriminating between VF and control samples in the discovery cohort using PepSeq Z scores. ROC curves for the 31 significantly enriched human peptides identified in the IgG screen (grey lines) and their sum (black), as well as ROC curves of PepSeq Z scores for the 16 peptides selected to be part of the peptide ELISA (orange) and the sum of this peptide set (red). AUCs indicated in lower right corner. **B)** Box plots showing the peptide ELISA normalized OD for discovery non-endemic control serum samples (light green, n=55), discovery VF patient sera that were run with PepSeq (dark green, n=51), validation non-endemic control (light purple, n=30), validation endemic control (medium purple, n=32), and validation VF patient sera (dark purple, n=78). Mann-Whitney U tests show both the VF PepSeq and VF independent have significantly increased optical density values (pvals < 0.001) compared to non-endemic (discovery and validation) and endemic (validation) controls. Horizontal dashed line shows the optimal OD value calculated based on the Discovery cohort (OD=0.092). **C)** ROC curves for peptide ELISA normalized ODs for VF samples run on PepSeq (Discovery, green), and independent validation VF samples run using all validation controls (dark purple), only endemic validation controls (medium purple) and only non-endemic validation controls (light purple).

### Characteristics of immunogenic proteins

Of the six proteins that were identified as showing significantly higher IgG reactivity in our VF cohort, only one has been previously identified as an immunogenic protein: spherical outer wall glycoprotein (SOWgp). Interestingly, SOWgp and IgG antigen 1 appear to be unique to *Coccidioides* spp., while the other four proteins have moderate sequence similarity (53-73% AA-level identity, 50-100% query coverage) to proteins from other fungi. To better understand the biological functions and immune accessibility of the six proteins identified, we utilized the AA sequence based prediction algorithms, deepLOC2.0 and SignalP-6.0, to predict the presence of transmembrane domains, cellular localization, and secretion signal peptides, (Supplemental Table 4)^32^. Four of the six proteins (SOWgp, IgG antigens 1-3) were predicted to be localized extracellularly with an average model confidence of 0.83. IgG antigen 1 was predicted to contain a secretion signal peptide and to be soluble (avg. confidence=0.90). Interestingly, the significantly enriched peptides for IgG antigen 2 were designed from domains that are shared between the two isoforms of this protein, both are predicted to have secretion signal peptides (model confidence = 0.99), however one isoform (KMP00029.1) was predicted to be soluble, whereas the other isoform (TPX26831.1) was predicted to be lipid anchored (model confidence = 0.84 and 0.96 respectively). SOWgp and IgG antigen 3 were both predicted to be lipid anchored with high confidence (0.94 and 0.98, respectively). IgG antigens 4 and 5 were predicted to be soluble (model confidence = 0.69 and 0.75, respectively), however, IgG antigen 4 was predicted to be localized to the cytoplasm (model confidence = 0.77) and IgG antigen 5 to the nucleus (model confidence = 0.77) (Supplemental Table 4). Additionally, transcriptional evidence suggests that four of the six proteins (SOWgp, IgG antigens 1-3) have increased gene expression during the host specific lifecycle (Table 1)^34,40^.

### Diagnostic potential of identified peptides

To assess the diagnostic potential of the immunogenic peptides within our discovery cohort, we generated receiver operating characteristic (ROC) curves for the validation sample cohort by ELISA. The area under the curve (AUC) for the 31 IgG peptides of interest individually ranged from 0.51 to 0.84, with an AUC of 0.88 when combining all 31 peptides using summed Z scores (Figure 3A). Using preliminary PepSeq data from an initial subset of our final dataset (n=101), we chose 16 peptides (from seven proteins) to include in a multipeptide ELISA (Supplemental Table 3). We generated ROC curves for each of the 16 peptides individually and combined, which resulted in a combined AUC of 0.89 (Figure 3A). Next, to validate the results from our highly-multiplexed assay in a simpler assay format, we created a multipeptide ELISA with these 16 peptides and used it to measure IgG antibody responses within a subset of our discovery cohort, as well as a validation cohort (new samples not assayed with PepSeq). From our discovery cohort, we randomly selected 51 VF and 55 non-endemic control samples to be assayed using the multipeptide ELISA. Our independent validation cohort consisted of 78 VF positive samples, and 62 control samples (32 endemic and 30 non-endemic). The VF positive samples had significantly higher ODs than the controls in both the discovery and validation cohorts (Control-discovery median=-0.02, VF-discovery median=0.93, Mann-Whitney pval= < 0.001) (Control-validation median=-0.01 VF-validation median=1.03, Mann-Whitney pval= < 0.001) (Figure 3C). ROC curves were used to assess the diagnostic potential of the ELISA. AUCs were 0.93 for the discovery cohort, 0.91 for the validation cohort (Figure 3D). When setting an optimal OD threshold (OD with maximum sensitivity + specificity) based on the discovery cohort data (OD=0.092), the sensitivity and specificity were 88% and 77% for the full validation cohort. Using the optimal OD 69/78 known positives were considered positive and 47/62 known negative controls were categorized negative.

### Detection of fungal antibodies using a commercial assay

In the human IgG PepSeq dataset, there were four samples from the discovery non-endemic control group that demonstrated reactivity to two or more of the 31 VF IgG enriched peptides. Additionally, in the ELISA assay, there were three non-endemic validation controls that had slightly elevated OD’s. To look for evidence of a past fungal infection, we tested the serum from these samples, along with three VF positives that showed high levels of IgG reactivity in our PepSeq assay, using commercially available reagents designed to detect antibody reactivity to *Coccidioides, Histoplasma, Aspergillus* and *Blastomyces* antigens. None of the discovery non-endemic control samples had detectable reactivity to any antigens tested and two out of three VF validation positives were reactive to the *Coccidioides* spp. antigen only. Because of volume limitations only two of the three non-endemic validation controls with elevated OD’s were assayed against the *Coccidioides* spp. antigen, both of which were negative (Supplemental Table 5).

## Discussion

In this study, we utilized highly-multiplexed serology to identify novel sero-reactive antigens from across the *Coccidioides* spp. proteome. Using a 244,000-plex *Coccidioides* specific PepSeq library (CWP), we probed VF positive and control sera from humans to determine reactivity profiles for IgG, IgA and IgM, and IgG in PTM. Human IgG samples had the highest amount of significantly reactive peptides, prompting the design of a pilot multi-peptide ELISA diagnostic and therefore, in this manuscript we primarily focused on antigens recognized by IgG antibodies.

We identified significantly higher IgG reactivity to 31 peptides from six different proteins in VF patients compared to controls. We also observed a significant positive correlation between reactive viral peptide counts and the summed Z scores against the 31 significantly enriched peptides within our VF cohort. This may indicate that there are differences in overall sample reactivity/quality within our VF cohort, which may explain why some of our VF discovery samples exhibited low reactivity against our 31 peptides of interest (Pearson correlation=0.353, pval=0.017) (Supplemental Figure 6).

Out of the 31 IgG peptides of interest, four peptides were from the spherule outer wall glycoprotein (SOWgp), a well characterized immunoreactive protein. This protein has previously been shown to be recognized by the cellular and humoral immune response to VF^41^. SOWgp is found on the cell walls of spherules and endospores, and is expressed primarily during the parasitic phase within the host^42^. It has been suggested that high titers of antibodies against SOWgp contribute to fungal cell opsonization and phagocytosis^43,44^. Additionally, antibody reactivity to SOWgp has been observed in VF positive patients, although this protein has not been utilized in diagnostic tests for VF^42–44^. Before our study, the epitope regions associated with antibody binding to SOWgp were not defined. Here, we highlight two linear regions commonly targeted by antibodies in patients with VF. We identified four significantly reactive peptides for SOWgp (CWP_085538, CWP_085539, CWP_103986, CWP_103987), three of which share an eight AA motif (“YCDTYGCD”) and likely contain a single epitope. The fourth peptide (CWP_085539) is designed from a different region of SOWgp and represents a second reactive epitope.

The remaining 27 peptides of interest come from five previously uncharacterized proteins. We used predictive algorithms to give insight into the potential biological activity and cellular localization for all six identified proteins. Four out of six of the identified proteins are predicted to be extracellular, which supports an interaction between the proteins and the immune system. In the literature we found limited studies discussing these uncharacterized proteins beyond those focused on transcript regulation, except for IgG antigen 1^34,40^. IgG antigen 1 is highly specific to *Coccidioides* spp. based on our BLASTP search and has been previously detected in *Coccidioides* cultures and later shown to be immunoreactive in dogs with VF^21,45^. Seven of the significantly reactive peptides in IgG antigen 1 were designed from amino acid positions 201-385, a region of the protein that is highly repetitive and proline rich. These peptides all include the same putative epitope, “PESKPFDPY”, where the bold and underlined letters indicate AAs conserved across all seven peptides, and the non-bold AAs are the consensus AAs (Supplemental Figure 8). Furthermore, two of the peptides contain two copies of this epitope (CWP_028597, CWP_054481), and these two peptides have the highest median Z score in the VF samples of any of the identified peptides of interest. Apart from this repeated epitope, IgG antigen 1 also has multiple other immunoreactive epitopes (Figure 1C,D), clearly indicating that it is a major target of the IgG antibody response to infections with *Coccidioides*.

A surprising result of our analysis was the lack of reactivity to any of the 45 peptides covering the current diagnostic IgG target, CF antigen, for which at least 48 VF patients had a positive CF titer (for 22 VF patients CF titer information was not available) (Supplemental Figures 4,5, Supplemental Table 6). One limitation of our approach is that the 30AA peptides can not accurately represent most conformational epitopes and they do not include post-translational modifications that might be present in the full antigens, *in vivo*. However, a GLM analysis showed a significant positive correlation between the summed Z score of our 31 IgG peptides of interest and CF titer, when controlling for gender, diagnosis, sample location, year of diagnosis and viral peptide counts (estimation of sample quality). Suggesting a positive relationship between CF titer and reactivity to *Coccidioides* peptides, with the exception of the four samples with the highest CF titers (1:256 and >1:1024), which showed very low PepSeq Z scores (Supplemental Figure 5). However, more samples are needed at these high CF titers to determine if the trends seen here are generalizable. Most VF patients develop a detectable IgG titer to CF during the first 1-3 months of infection. The titer slowly decreases as disease resolves, with an apparent half-life of ~90 days for a doubling dilution ^2,46,47^. Interestingly, we were able to detect IgG antibody reactivity to the 31 peptides of interest in samples collected years after diagnosis (Supplemental Figure 7) indicating the presence of long lived antibodies in response to *Coccidioides* spp. infection.

To validate the IgG reactivity results from PepSeq, a subset of the 31 highly reactive peptides were commercially synthesized as 30mers (without the additional constant region amino acids and DNA tag present in the PepSeq library) and used to build a multi-epitope ELISA. The selection was based on preliminary data analysis and includes nine different proteins (including peptides from all six significant proteins of interest described here), with one to four peptide sequences per protein (Supplemental Table 3). We tested the ability of the ELISA assay to discriminate between a Validation cohort consisting of VF patients, more blood bank controls and VF negative samples from the endemic region all of which had not previously been assayed with our CWP PepSeq library and thus did not inform peptide selection. Using the independent validation sample set, we demonstrated the peptides identified in this study could successfully discriminate between VF patients and blood bank controls/VF negatives at high accuracy (ROC 0.91). These data support that binding signals to these linear epitopes, in an ELISA format, can differentiate between VF positive and negative donors with sensitivity (88%) and specificity (77%) on par with current clinical diagnostics^5,13,18,48^.

An asset to our design is the control samples from the endemic region are from patients that were screened for coccidioidomycosis during differential diagnosis or prior to transplant. All of these patients tested negative for *Coccidioides* infection using current clinical assays (CF, ID and EIA). In general, these patients had higher ODs compared to the blood bank control samples collected outside of the endemic region. This could be due to previous exposure or infection with *Coccidioides*, which was resolved prior to testing. Two of these patients had OD values in our peptide ELISA that would be considered positive based on the optimal OD threshold. Interestingly, these patients exhibit strong reactivity to other *Coccidioides* antigens of interest measured by NAPPA and MISPA assays ^49^. Chart review of these patients revealed one patient had previously been VF positive by EIA a year prior and the other did not appear to have background that would support a current VF diagnosis. We propose these patients may recognize other *Coccidioides* antigens and may could be false positives as they do not demonstrate antibody reactivity to chitinase 1 (CTS1), the antigen used in both clinical tests, further highlighting the need for additional diagnostic targets beyond CTS1.

Although our sample size is modest, the statistical approach in PepSeq utilized here is robust with zero peptides being detected as significantly enriched in the IgG non-endemic control cohort when running a flipped analysis. Furthermore, our random permutation analysis of the human IgG data indicated that our false discovery rate was quite low. It is likely that there are more reactive peptides and proteins of interest that we are not powered to identify as significant with our current cohort sizes, or that require secondary or tertiary structure for binding. The VF discovery samples utilized in PepSeq assays were originally collected for a study focused on long lived cellular immunity and thus have a wide range of days from diagnosis to collection. Having such a broad sampling range of time from diagnosis may have negatively impacted our ability to detect disease specific (early v late) antibodies, with the relative abundance of *Coccidioides* spp. specific antibodies waning over time. However, having a diversity of samples also increased the disease space covered in this study and may have revealed antigens associated across disease states. Another limitation to consider is that the discovery control cohorts in the PepSeq study were obtained from a blood bank in Louisiana, and we have no metadata for these samples other than where the collection was made. While these samples were taken from outside the endemic VF region, it is possible that some of them previously lived in or visited a *Coccidioides* spp. endemic region where they were exposed to the fungus. This is one possible explanation for some of the reactivity we see in the control cohort samples in Figure 1C. However, using a commercial immunodiffusion kit, we did not find evidence of antibody reactivity against *Coccidioides* spp. or the other endemic fungi on the panel. Of note, we also only observed two out of three confirmed VF patients show reactivity in the immunodiffusion assay run with the same commercially available reagents. The VF patient that tested negative was tested at the UC Davis lab, which uses different sample preparation procedures and different antigen lysates. At UC Davis, the sample was negative by CF testing, and positive by immunodiffusion testing.

In addition to our IgG investigations, we measured IgA and IgM reactivity in a subset of the discovery cohort. We identified two peptides with significantly increased IgA reactivity in VF patients compared to non-endemic controls (Supplemental Figure 2B,C). Interestingly, while not significant, at the cohort level, we did observe a small number of VF samples showing elevated IgA reactivity to 16 of the peptides identified as significant in the IgG screen (Supplemental Figure 3C). These results suggest that, in addition to IgG, humans also produce circulating IgA antibodies in response to *Coccidioides* spp. infection. Comparing the IgG and IgA reactivity profiles for the two significant peptides (CWP_054480, CWP_054481), there appears to be less specificity with IgA assays compared to the IgG assays. We did not identify any peptides with significantly elevated IgM reactivity in our VF cohort, and only observed modest reactivity in 1-2 samples to 6 of the peptides of interest determined from our IgG screen (Supplemental Figure 3B).

Lastly, we report the first in-depth antigenic analysis of IgG antibody reactivity against *Coccidioides* spp. in naturally infected PTMs. We detected significantly higher reactivity in our animals with VF to two peptides designed from the IgG antigen 3 protein that was also identified in our human IgG PepSeq screen (described above). Interestingly, among the 31 significant peptides identified in the human IgG PepSeq screen, a subset of naturally infected PTMs exhibited high levels of reactivity towards antigens commonly targeted in human CM patients (Supplemental Figure 3D). PTM are being used in *Coccidioides* spp. vaccine trials and a better understanding of the B cell response to natural infection in these animals may be valuable in these efforts. While our sample size is limited, our results show that PepSeq can be used to detect antibody reactivity in PTMs and that B cell responses to infection in humans and PTMs target many of the same proteins/epitopes, though not necessarily as consistently. Future studies will be needed to further map the B cell responses in PTMs in response to natural infection with *Coccidioides* spp.

## Funding

This work was supported by 5R21AI180987-02 (Mead), U19 AI166058-01 (Keim) and Developmental Research Award under U19 AI166058-01 (Elko). WaNPRC is supported by grant P51 OD010425 and U42OD011123. The funders did not play any role in the study design, data collection and analysis, decision to publish, or preparation of the manuscript

### Patent pending

Compositions and Methods for Detection and Treatment of Coccidioides Infection to Heather Lynn Mead, John Altin, Jason Ladner, Evan Elko.

---

**Supplemental Figure 1.**
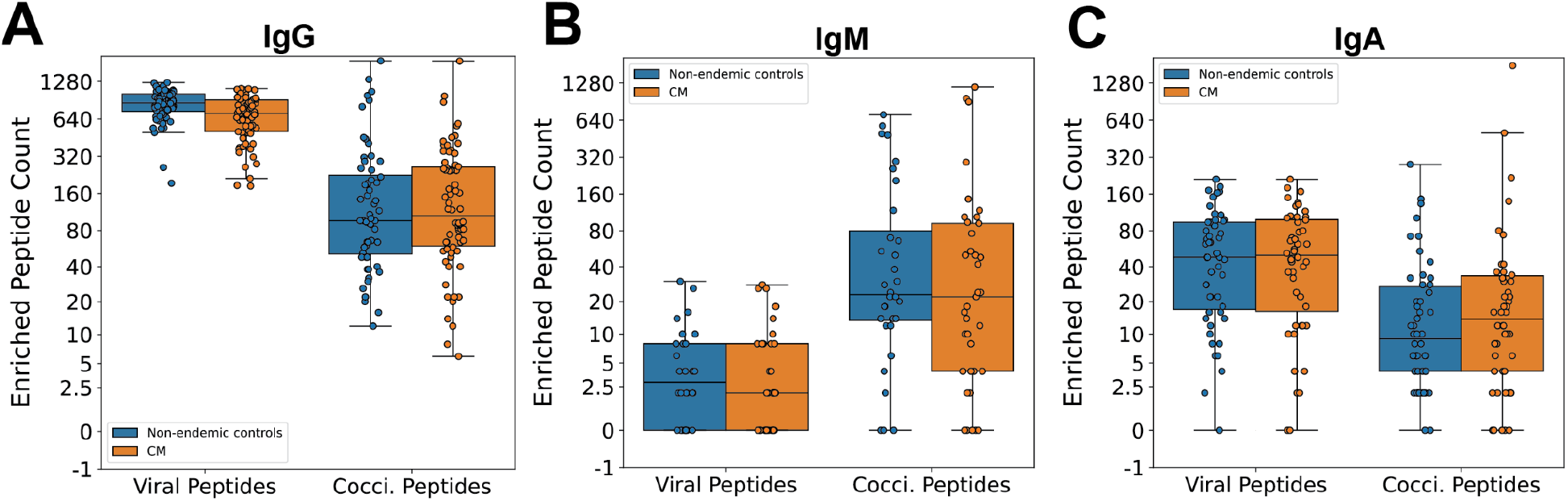
Enriched peptides in VF and control cohorts. Box plots show the number of enriched viral positive control and *Coccidioides* derived peptides (Z ≥ 10 in both replicates) in the discovery negative control and VF cohorts for human IgG (A), human IgM (B) and human IgA (C).

**Supplemental Figure 2.**
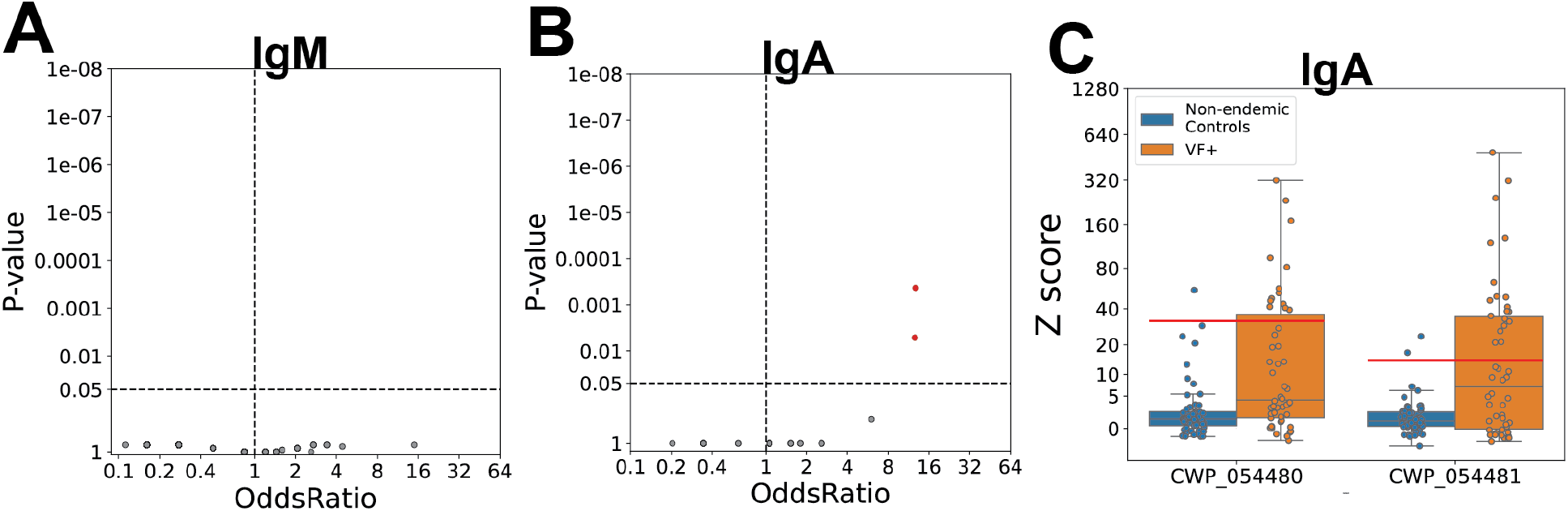
PepSeq analysis of IgM and IgA. **A**) A volcano plot showing the −log10(P-value) from Fisher’s exact test versus log10(OddsRatio) for a subset of 273 peptides recognized by serum IgM antibodies in ≥5 serum samples. Horizontal dashed line indicates a Benjamini-Hochberg corrected P-value of 0.05. Vertical dashed line indicates an odds ratio of 1. **B**) A volcano plot showing the −log10(P-value) from a Fisher’s exact test versus log10(OddsRatio) for a subset of 32 peptides recognized by serum IgA antibodies in ≥5 serum samples. Horizontal dashed line indicates a Benjamini-Hochberg corrected P-value of 0.05. Vertical dashed line indicates an odds ratio of 1. **C)** Boxplots showing the Z score distribution of the two significantly enriched IgA peptides in the non-endemic control and VF cohorts.

**Supplemental Figure 3.**
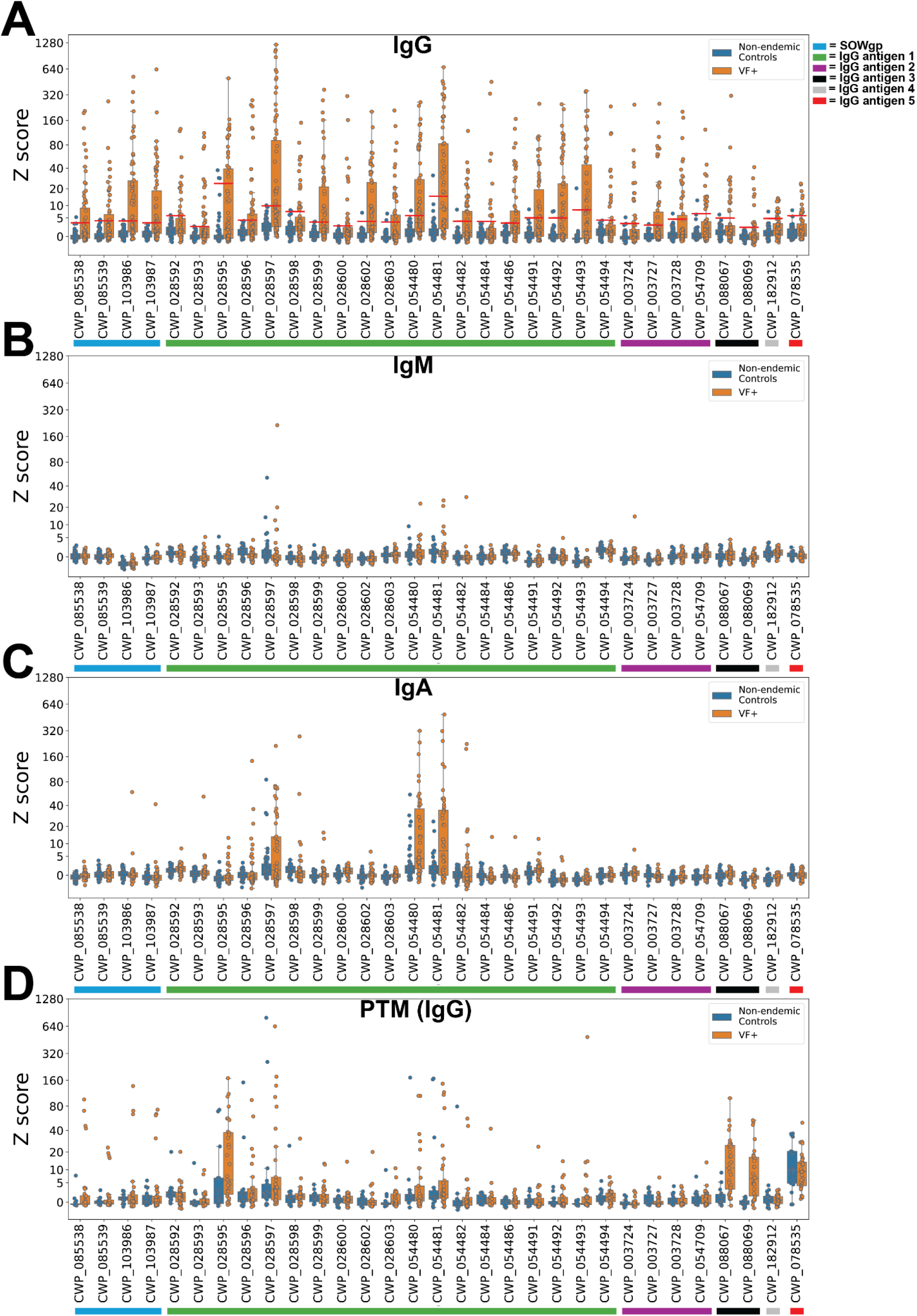
Reactivity profiles for 31 IgG peptides of interest for human IgM, human IgA and PTM IgG. Box plots show the Z scores for the 31 peptides of interest identified in the human IgG screen (A), in human IgM (B), human IgA (C), and PTM IgG (D).

**Supplemental Figure 4.**
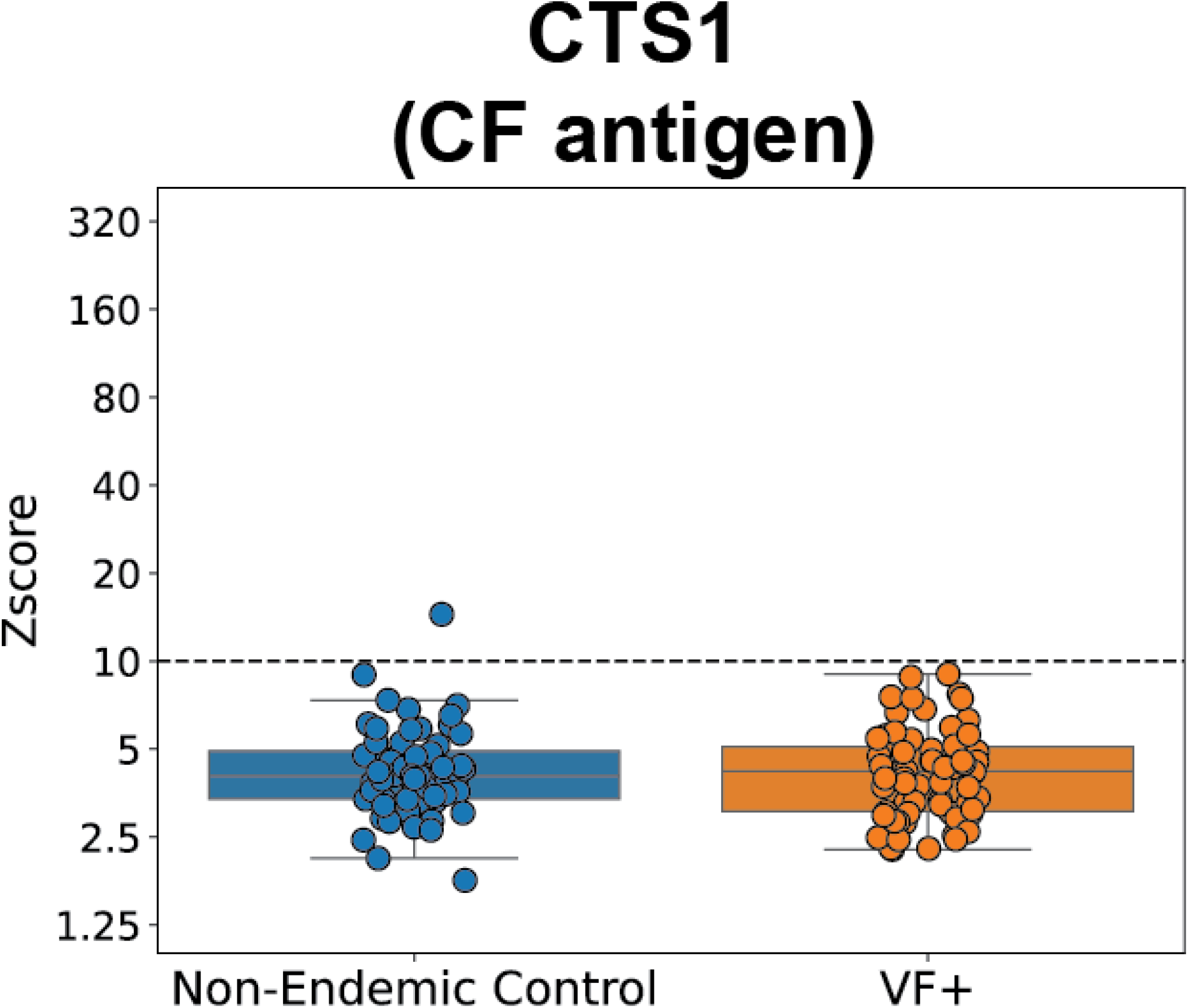
PepSeq IgG reactivity for CF antigen (CTS1). Box plots showing the Z score distributions of the max Z score for all peptides (n=45) designed from CF antigen (CTS1).

**Supplemental Figure 5.**
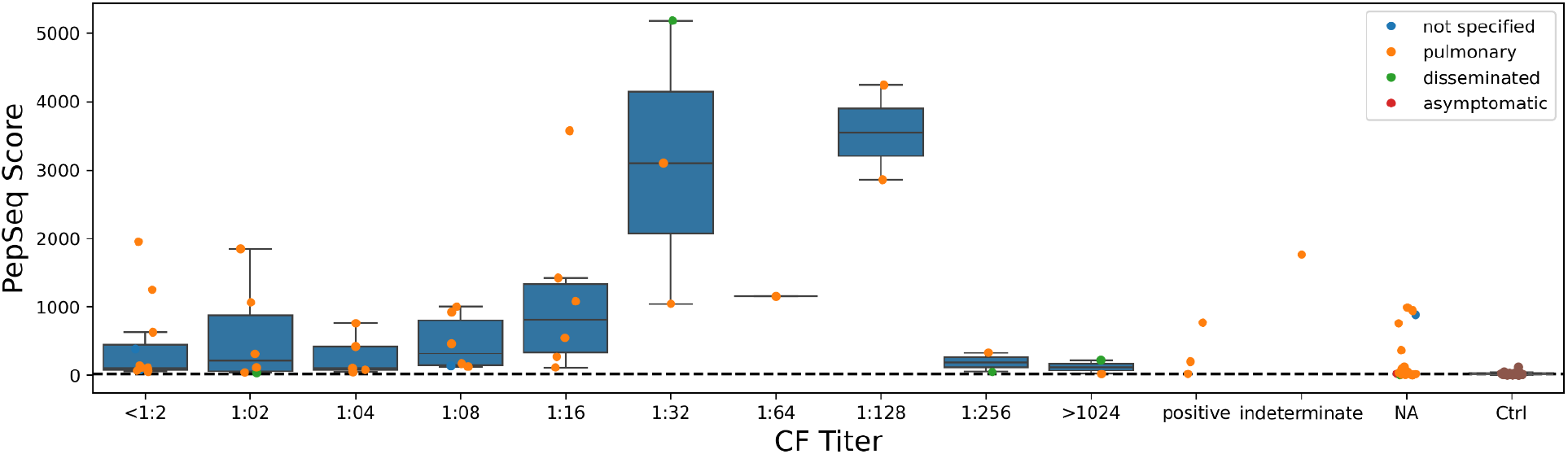
Correlation of PepSeq score and CF Titer. Box plots showing the sum of the IgG Z scores for the 31 IgG peptides of interest for each clinically measured CF titer. The color of the points indicate the diagnosis of the patients if available.

**Supplemental Figure 6.**
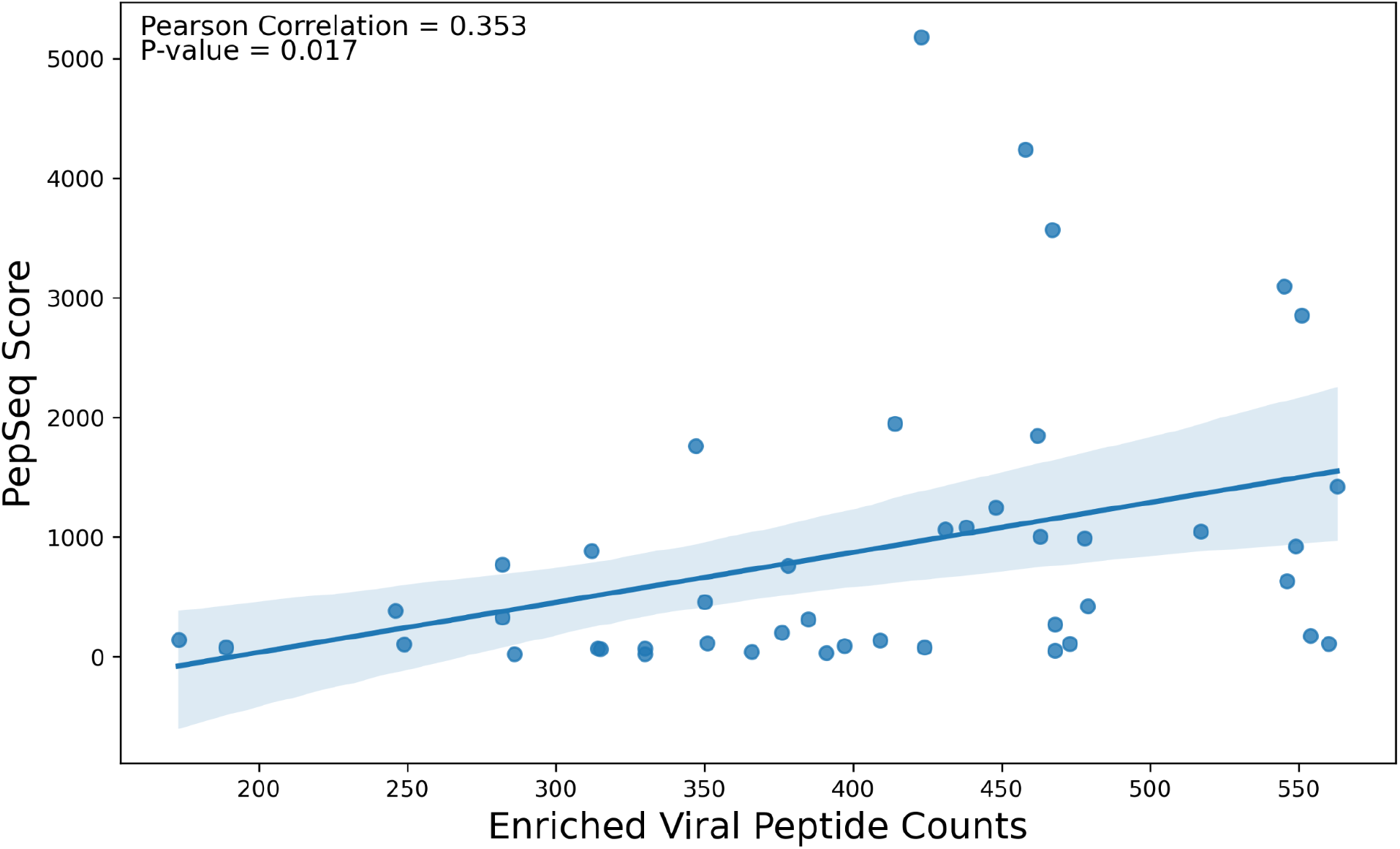
Correlation of PepSeq scores for the 31 IgG peptides of interest and the counts of enriched viral peptides for each VF discovery cohort sample. Scatter plot showing the sum of the IgG Z scores for the 31 IgG peptides of interest (y-axis) and the number of control, viral enriched peptides (x-axis). Pearson correlation coefficient is 0.353 with a p-value of 0.017. The blue line shows the linear regression and the shaded regions show the 95% confidence interval.

**Supplemental Figure 7.**
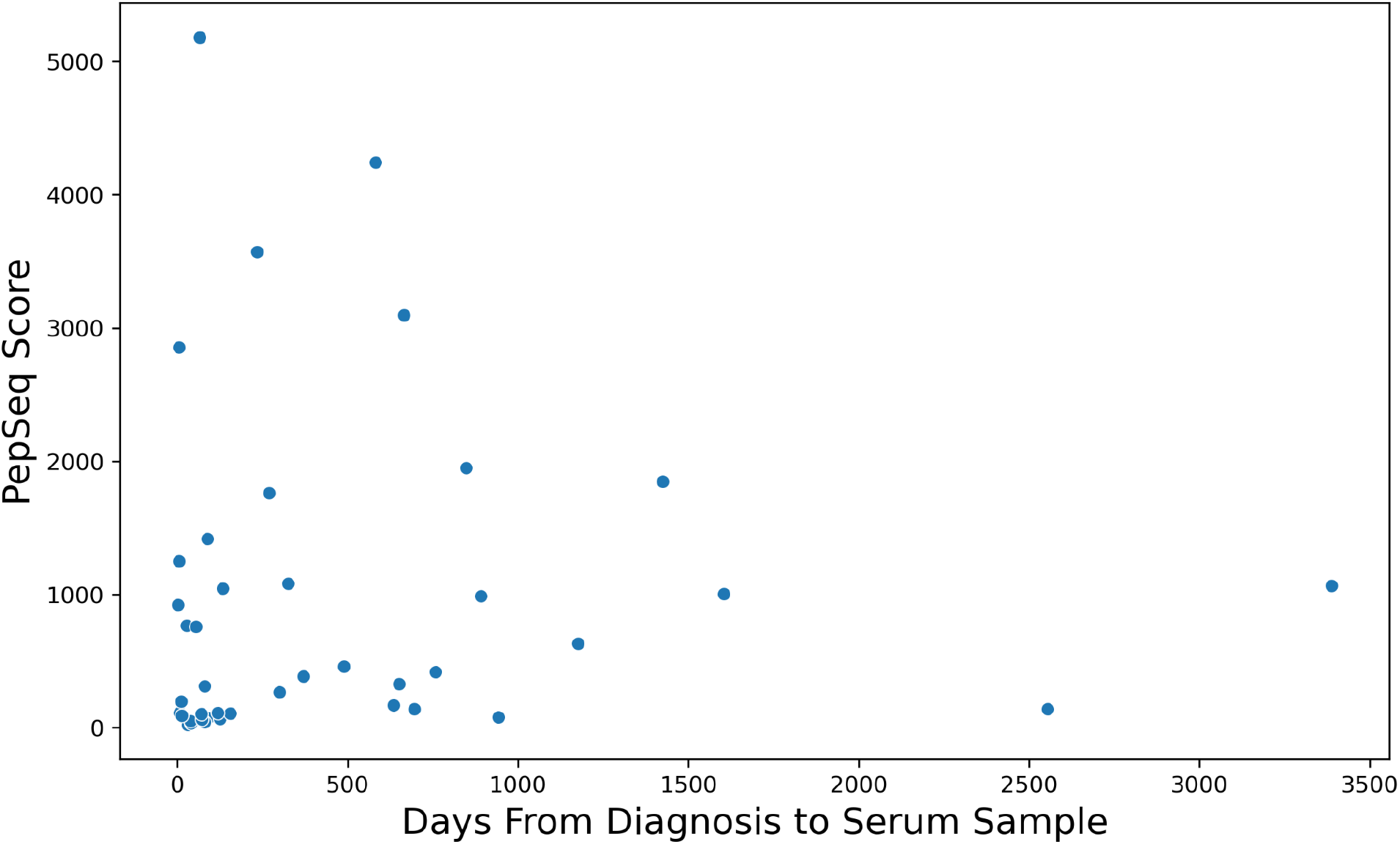
Correlation of PepSeq scores for the 31 IgG peptides of interest and sample collection time from diagnosis. Scatter plot showing the sum of the IgG Z scores for the 31 IgG peptides of interest (y-axis) and the days between when the sample was collected and VF diagnosis (x-axis). Antibody reactivity is detected years after diagnosis VF patient samples were collected 0-3,388 days after diagnosis.

**Supplemental Figure 8.**
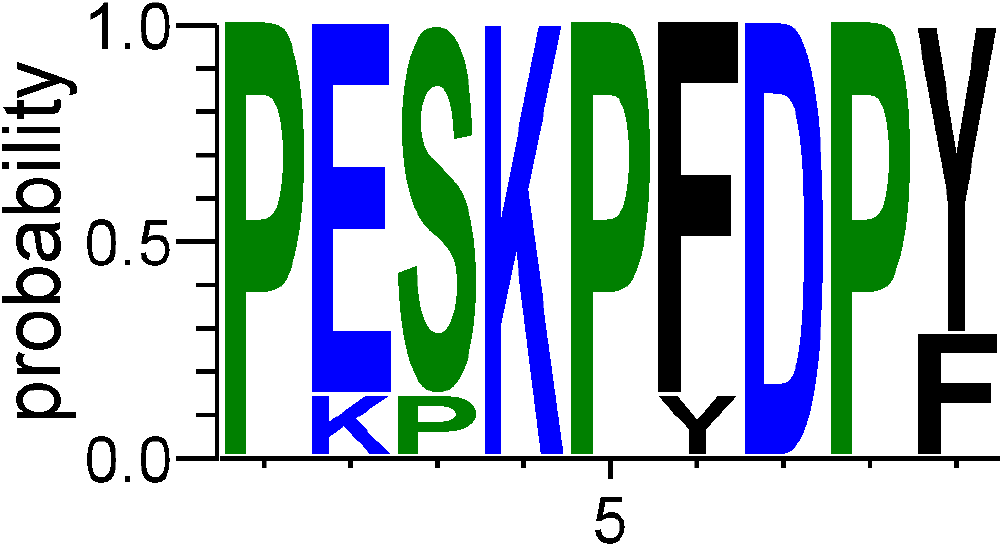
Sequence conservation of reactive epitope found in IgG antigen 1. Logo plot showing the AA probability at each position of the highly reactive repetitive proline rich epitope identified in IgG antigen 1.

## References

1. Bays, D. J. & Thompson, G. R., 3rd. Coccidioidomycosis. Infect. Dis. Clin. North Am. 35, 453–469 (2021).

2. Gabe, L. M., Malo, J. & Knox, K. S. Diagnosis and Management of Coccidioidomycosis. Clin. Chest Med. 38, 417–433 (2017).

3. Galgiani, J. N. et al. 2016 Infectious Diseases Society of America (IDSA) Clinical Practice Guideline for the Treatment of Coccidioidomycosis. Clin. Infect. Dis. 63, e112–46 (2016).

4. Donovan, F. M. et al. Delays in coccidioidomycosis diagnosis and associated healthcare utilization, Tucson, Arizona, USA. Emerg. Infect. Dis. 25, 1745–1747 (2019).

5. Theel, E. S. et al. Practical Guidance for Clinical Microbiology Laboratories: Antibody and antigen detection methods for dimorphic fungal infections. Clin. Microbiol. Rev. 38, e0000520 (2025).

6. McHardy, I. H., Barker, B. & Thompson, G. R., 3rd. Review of Clinical and Laboratory Diagnostics for Coccidioidomycosis. J. Clin. Microbiol. 61, e0158122 (2023).

7. Hung, C. Y., Yu, J. J., Lehmann, P. F. & Cole, G. T. Cloning and expression of the gene which encodes a tube precipitin antigen and wall-associated beta-glucosidase of Coccidioides immitis. Infect. Immun. 69, 2211–2222 (2001).

8. Cox, R. A. & Britt, L. A. Antigenic identity of biologically active antigens in coccidioidin and spherulin. Infect. Immun. 55, 2590–2596 (1987).

9. Johnson, S. M. & Pappagianis, D. The coccidioidal complement fixation and immunodiffusion-complement fixation antigen is a chitinase. Infect. Immun. 60, 2588–2592 (1992).

10. Durkin, M. et al. Diagnosis of coccidioidomycosis with use of the Coccidioides antigen enzyme immunoassay. Clin. Infect. Dis. 47, e69–73 (2008).

11. Kassis, C. et al. Diagnosis of Coccidioidomycosis with the Second-Generation Miravista IgG and IgM Enzyme Immunoassay and the Role of Adding Miravista Coccidioides Antigen Detection to Immunodiagnostic Assays. Med. Mycol. 62, (2024).

12. Grill, F. J. et al. Clinical Laboratory Utility of a Humanized Antibody in Commercially Available Enzyme Immunoassays for Coccidioidomycosis. Microbiol Spectr e0257322 (2022) doi:10.1128/spectrum.02573-22.

13. Grill, F. J. et al. Development of a rapid lateral flow assay for detection of anti-coccidioidal antibodies. J. Clin. Microbiol. 61, e0063123 (2023).

14. Pappagianis, D. & Zimmer, B. L. Serology of coccidioidomycosis. Clin. Microbiol. Rev. 3, 247–268 (1990).

15. Khan, S. et al. Interlaboratory agreement of coccidioidomycosis enzyme immunoassay from two different manufacturers. Med. Mycol. 57, 441–446 (2019).

16. Donovan, F. M. et al. Comparison of a Novel Rapid Lateral Flow Assay to Enzyme Immunoassay Results for Early Diagnosis of Coccidioidomycosis. Clin. Infect. Dis. 73, e2746–e2753 (2021).

17. Stevens, D. A. et al. Comparative study of newer and established methods of diagnosing coccidioidal meningitis. J. Fungi (Basel) 6, 125 (2020).

18. Malo, J. et al. Comparison of three anti-coccidioides antibody enzyme immunoassay kits for the diagnosis of coccidioidomycosis. Med. Mycol. 58, 774–778 (2020).

19. Barker, B. M., Jewell, K. A., Kroken, S. & Orbach, M. J. The population biology of coccidioides: epidemiologic implications for disease outbreaks. Ann. N. Y. Acad. Sci. 1111, 147–163 (2007).

20. Neafsey, D. E. et al. Population genomic sequencing of Coccidioides fungi reveals recent hybridization and transposon control. Genome Res. 20, 938–946 (2010).

21. Mitchell, N. M. et al. Proteogenomic re-annotation of Coccidioides posadasii strain Silveira. Proteomics 18, (2018).

22. Lequin, R. M. Enzyme immunoassay (EIA)/enzyme-linked immunosorbent assay (ELISA). Clin Chem 51, 2415–2418 (2005).

23. Henson, S. N. et al. PepSeq: a fully in vitro platform for highly multiplexed serology using customizable DNA-barcoded peptide libraries. Nat. Protoc. https://doi.org/10.1038/s41596-022-00766-8 (2022) doi:10.1038/s41596-022-00766-8.

24. Piña, A. et al. Mapping disparities in viral infection rates using highly multiplexed serology. mSphere 9, e0012724 (2024).

25. Elko, E. A. et al. Recurrent SARS-CoV-2 mutations at Spike D796 evade antibodies from pre-Omicron convalescent and vaccinated subjects. Microbiol Spectr e0329123 (2024) doi:10.1128/spectrum.03291-23.

26. Elko, E. A. et al. COVID-19 vaccination elicits an evolving, cross-reactive antibody response to epitopes conserved with endemic coronavirus spike proteins. Cell Rep. 40, 111022 (2022).

27. Kelley, E. J. et al. Virome-wide detection of natural infection events and the associated antibody dynamics using longitudinal highly-multiplexed serology. Nat. Commun. 14, 1783 (2023).

28. Ladner, J. T. et al. Epitope-resolved profiling of the SARS-CoV-2 antibody response identifies cross-reactivity with endemic human coronaviruses. Cell Rep Med 2, 100189 (2021).

29. Elko, E. A. et al. PepSeq as a highly multiplexed platform for melioidosis antigen discovery and vaccine development. Front. Immunol. 16, (2025).

30. Vita, R. et al. The Immune Epitope Database (IEDB): 2024 update. Nucleic Acids Res. 53, D436–D443 (2025).

31. Edgar, R. C. Search and clustering orders of magnitude faster than BLAST. Bioinformatics 26, 2460–2461 (2010).

32. Thumuluri, V., Almagro Armenteros, J. J., Johansen, A. R., Nielsen, H. & Winther, O. DeepLoc 2.0: multi-label subcellular localization prediction using protein language models. Nucleic Acids Res. 50, W228– W234 (2022).

33. Teufel, F. et al. SignalP 6.0 predicts all five types of signal peptides using protein language models. Nat. Biotechnol. 40, 1023–1025 (2022).

34. Basenko, E. Y. et al. FungiDB: An Integrated Bioinformatic Resource for Fungi and Oomycetes. J Fungi (Basel) 4, (2018).

35. Sayers, E. W. et al. Database resources of the National Center for Biotechnology Information in 2025. Nucleic Acids Res. 53, D20–D29 (2025).

36. Seabold, S. & Perktold, J. Statsmodels: Econometric and statistical modeling with python. in Proceedings of the 9th Python in Science Conference (SciPy, 2010). doi:10.25080/majora-92bf1922-011.

37. Sievers, F. et al. Fast, scalable generation of high-quality protein multiple sequence alignments using Clustal Omega. Mol. Syst. Biol. 7, 539 (2011).

38. Guerriero, K. A. et al. Recrudescence of natural coccidioidomycosis during combination antiretroviral therapy in a pigtail macaque experimentally infected with simian immunodeficiency virus. AIDS Res. Hum. Retroviruses 37, 505–509 (2021).

39. Koistinen, K. et al. Coccidioidomycosis in nonhuman primates: Pathologic and clinical findings. Vet. Pathol. 55, 905–915 (2018).

40. Whiston, E. et al. Comparative transcriptomics of the saprobic and parasitic growth phases in Coccidioides spp. PLoS One 7, e41034 (2012).

41. Hung, C. Y., Ampel, N. M., Christian, L., Seshan, K. R. & Cole, G. T. A major cell surface antigen of Coccidioides immitis which elicits both humoral and cellular immune responses. Infect. Immun. 68, 584–593 (2000).

42. Cole, G. T. et al. Isolation and morphology of an immunoreactive outer wall fraction produced by spherules of Coccidioides immitis. Infect. Immun. 56, 2686–2694 (1988).

43. Hung, C.-Y. et al. A metalloproteinase of Coccidioides posadasii contributes to evasion of host detection. Infect. Immun. 73, 6689–6703 (2005).

44. Cox, R. A. & Magee, D. M. Coccidioidomycosis: host response and vaccine development. Clin. Microbiol. Rev. 17, 804–39, table of contents (2004).

45. Koehler, M. A. et al. Discovery of a unique set of dog-seroreactive Coccidioides proteins using Nucleic Acid Programmable Protein Array. J. Fungi (Basel) 10, 307 (2024).

46. McHardy, I. H. et al. Coccidioidomycosis Complement Fixation Titer Trends in the Age of Antifungals. J. Clin. Microbiol. 56, (2018).

47. Smith, C. E., Saito, M. T. & Simons, S. A. Pattern of 39,500 serologic tests in coccidioidomycosis. J. Am. Med. Assoc. 160, 546–552 (1956).

48. Grys, T. E. et al. Comparison of two FDA-cleared EIA assays for the detection of Coccidioides antibodies against a composite clinical standard. Med. Mycol. 57, 595–600 (2019).

49. Mitchell Magee, D. High-throughput Antibody Immune Profiling of Valley Fever Patients. (2026).

